# Response of Bat Activity to Land-Cover and Land-Use Change in Savannas is Scale-, Season-, and Guild-Specific

**DOI:** 10.1101/613943

**Authors:** Julie Teresa Shapiro, Ara Monadjem, Timo Röder, Robert A. McCleery

## Abstract

Tropical savannas are biomes of global importance that are under severe pressure from anthropogenic change, including land-cover and land-use change. Bats, the second-most diverse group of mammals, are critical to ecosystem functioning, but may be vulnerable to such anthropogenic stresses. However, there is little information on the response of savanna bats to land-cover and land-use change, especially in Africa. This limits our ability to develop conservation strategies for bats and maintain the ecosystem functions and services they provide in this biome. Using acoustic monitoring, we measured how guild-specific (aerial, edge, and clutter forager) bat activity responded to both fine-scale metrics of vegetation structure and landscape-scale metrics of land-cover composition and configuration across the wet and dry seasons in a savanna in southern Africa undergoing rapid land-cover and land-use change. We found that all three guilds responded more strongly to landscape metrics than fine-scale vegetation structure, although the specific metrics varied between guilds. Aerial and edge bats responded most strongly to the percent savanna cover and savanna fragmentation in both seasons while clutter bats responded to percent rural cover in the wet season and percent water cover in the dry. All three guilds responded more strongly to the landscape in the dry season than the wet season. Our results show it is possible to conserve bats, and the ecosystem services they can provide, in savannas undergoing anthropogenic land-use and land-cover change but strategies to do so must consider foraging guild, large spatial scales, and seasonal variation in bat activity.

**Highlights:**

- Bats in savannas respond to land-cover and land-use change on large spatial scales
- Landscape had a greater influence on bat activity in the dry season than the wet
- Aerial and edge forager activity responded to savanna cover and fragmentation
- Clutter forager activity was best explained by rural and water cover
- Minimizing fragmentation and maintaining water promotes bat activity in modified savannas

## 1. Introduction

Tropical savannas are biomes of global importance for people and wildlife (Bond and Parr, 2010; Murphy et al., 2016; Parr et al., 2014). They contain high levels of biodiversity, provide essential habitat for endemic and endangered species (Murphy et al., 2016), account for a large amount of terrestrial net primary productivity, and store carbon (Parr et al., 2014). Savannas also provide essential resources to people, especially in developing countries, such as pasture for livestock, firewood, thatching materials, and medicinal plants (Egoh et al., 2009; Fensham et al., 2005; Hoffmann et al., 2012; Parr et al., 2014; van der Werf et al., 2010).

Despite their importance, tropical savannas are generally underappreciated, understudied and under-protected (Laurance et al., 2014; Parr et al., 2014), with less than 13% under any kind of official protection (Jenkins and Joppa, 2009). Globally, one of the principal threats to tropical savannas is land-cover change, particularly the conversion of savanna to agriculture, including both low-intensity croplands and high intensity commercial production (Aleman et al., 2016; Laurance et al., 2014).

Land-cover change has profound, often negative, impacts on wildlife (Foord et al., 2018; Reynolds et al., 2018; Sala et al., 2000). At fine spatial scales, land-cover change alters the type and structure of vegetation, eliminating foraging habitat or shelter (Fahrig et al., 2011; Goodwin et al., 2002; Tscharntke et al., 2012). On larger scales, landscape composition (the different types of land cover) and configuration (the spatial pattern of land cover) affect wildlife through different mechanisms. Changes in landscape composition typically lead to reductions in native habitats and the loss of resources located in them (Fischer and Lindenmayer, 2006; Tscharntke et al., 2012). In contrast, changes in landscape configuration, regardless of the amount of cover, affect wildlife through edge effects, patch isolation, and loss of connectivity across the landscape (Fahrig, 2003).

Bats in savannas appear to respond to land cover changes (Mtsetfwa et al., 2018; Weier et al., 2018) and may serve as bioindicators (Jones et al., 2009). They are the second most diverse order of mammals (Burgin et al., 2018) and provide important ecosystem services such as pest control, pollination, and seed dispersal (Boyles et al., 2011; Kunz et al., 2011; Maas et al., 2013; Taylor et al., 2017; Williams-Guillén et al., 2008). There is growing evidence that in savannas in particular, some bat species exhibit strong preferences for agricultural landscapes (Noer et al., 2012; Toffoli and Rughetti, 2017) where they play an important role in consuming pest insects (Bohmann et al., 2011; Puig-Montserrat et al., 2015; Taylor et al., 2013, 2018, 2017).

Conserving bats, and therefore maintaining the ecosystem services and functions that they provide, requires an understanding of how bats are affected by land-cover change and at what spatial scale these changes most affect them. Although we know that bats respond to changes in both fine-scale vegetation structure and landscape-scale composition and configuration (Brigham et al., 1997; Fuentes-Montemayor et al., 2013; Gehrt and Chelsvig, 2003; Kalda et al., 2015; Monadjem and Reside, 2008) these relationships are often a function of spatial scale (Gorresen et al., 2005; Mendes et al., 2017; Pinto and Keitt, 2008). Additionally, bats’ response to land cover varies greatly between regions, biomes, seasons (Ferreira et al., 2017; Klingbeil and Willig, 2010; Mendes et al., 2014), and species or guilds (Gorresen et al., 2005; Klingbeil and Willig, 2009; Mendes et al., 2017; Müller et al., 2012).

To date, most research on the impacts of land-cover change on bats has been conducted in forest biomes (Estrada-Villegas et al., 2010; Ferreira et al., 2017; Pinto and Keitt, 2008; Williams-Guillén and Perfecto, 2011), limiting our ability to generalize patterns. Our understanding of how land-cover change affects bats in savannas, particularly in Africa, is far more limited (Meyer et al., 2016; Monadjem and Reside, 2008; Mtsetfwa et al., 2018; Weier et al., 2018). There is evidence that high intensity agriculture in southern African savannas can negatively affect some bat species (Mtsetfwa et al., 2018), while remnant natural and semi-natural vegetation (Mtsetfwa et al., 2018; Weier et al., 2018) and wetlands (Sirami et al., 2013) in these landscapes can promote bat activity. However, the role of landscape configuration has not been considered. In addition, the relative effects of fine-scale vegetation compared to landscape composition and configuration have not been directly compared. Finally, studies in this region have only compared the effects of savanna and commercial agriculture on bats (Mtsetfwa et al., 2018; Sirami et al., 2013; Weier et al., 2018), while the role of rural areas and villages has been largely neglected, although they comprise a large, and growing, component of the landscape (Bailey et al., 2015).

In order to understand the effects of land-cover change on bats in tropical savannas, we measured guild-level responses in bat activity across the wet and dry seasons to both fine-scale metrics of vegetation structure and landscape-scale metrics of land cover composition and configuration across northeastern Eswatini (formerly Swaziland). This region is part of the Maputaland-Albany-Pondoland biodiversity hotspot (Steenkamp et al., 2005) and undergoing rapid land-cover change, primarily as a result of agricultural expansion and intensification (Bailey et al., 2015). Our objectives were to: 1) quantify the response of bats to variation in fine-scale vegetation structure and landscape-scale land-cover composition and configuration; 2) compare the variation in responses by foraging guild; 3) determine the most relevant spatial scale of the response for each guild; and 4) ascertain how responses vary by season. We expected to see guild-specific responses to both fine- and landscape-scale characteristics, with bats that use denser vegetation and fly shorter distances responding more strongly to fine-scale vegetation structure while bats that forage in open areas and fly longer distances were expected to respond more strongly to landscape-scale characteristics (Ferreira et al., 2017; Fuentes-Montemayor et al., 2013; Pinto and Keitt, 2008). In general, we expected to see a greater effect of landscape composition than configuration on bats, as has been reported in previous studies (Arroyo-Rodríguez et al., 2016; Meyer and Kalko, 2008). We also expected to see strong seasonal variation in response from all guilds (Monadjem and Reside, 2008; Mtsetfwa et al., 2018; Taylor et al., 2013).

## 2. Materials and Methods

### 2.1 Study Area

This study was conducted across an area of approximately 2,300 km^2^ in the eastern low-lying region of Eswatini referred to as the “Lowveld” which is bordered by the Drakensberg Mountains in the west and the Lubombo Mountains in the east (Figure 1). The area is a part of the Maputaland-Pondoland-Albany biodiversity hotspot (Steenkamp et al., 2005), which stretches from southern Mozambique, through eastern Eswatini, and into South Africa. This region has been subject to rapid land-cover change, mainly from expansion of commercial and small-holder croplands (Bailey et al., 2015). Elevation ranges from approximately 150 m to 600 m above sea level. The Lowveld is characterized by a warm, semi-arid subtropical climate (Matondo et al. 2004). The annual mean temperature is 20-22° C, with a mean monthly temperature of 26° C in January and 18° C in July (Monadjem and Garcelon, 2005). Annual rainfall is 500-700 mm per year, concentrated in the summer months of October to March (Matondo et al. 2004; Monadjem and Reside 2008; Knox et al. 2010).

**Figure 1.**
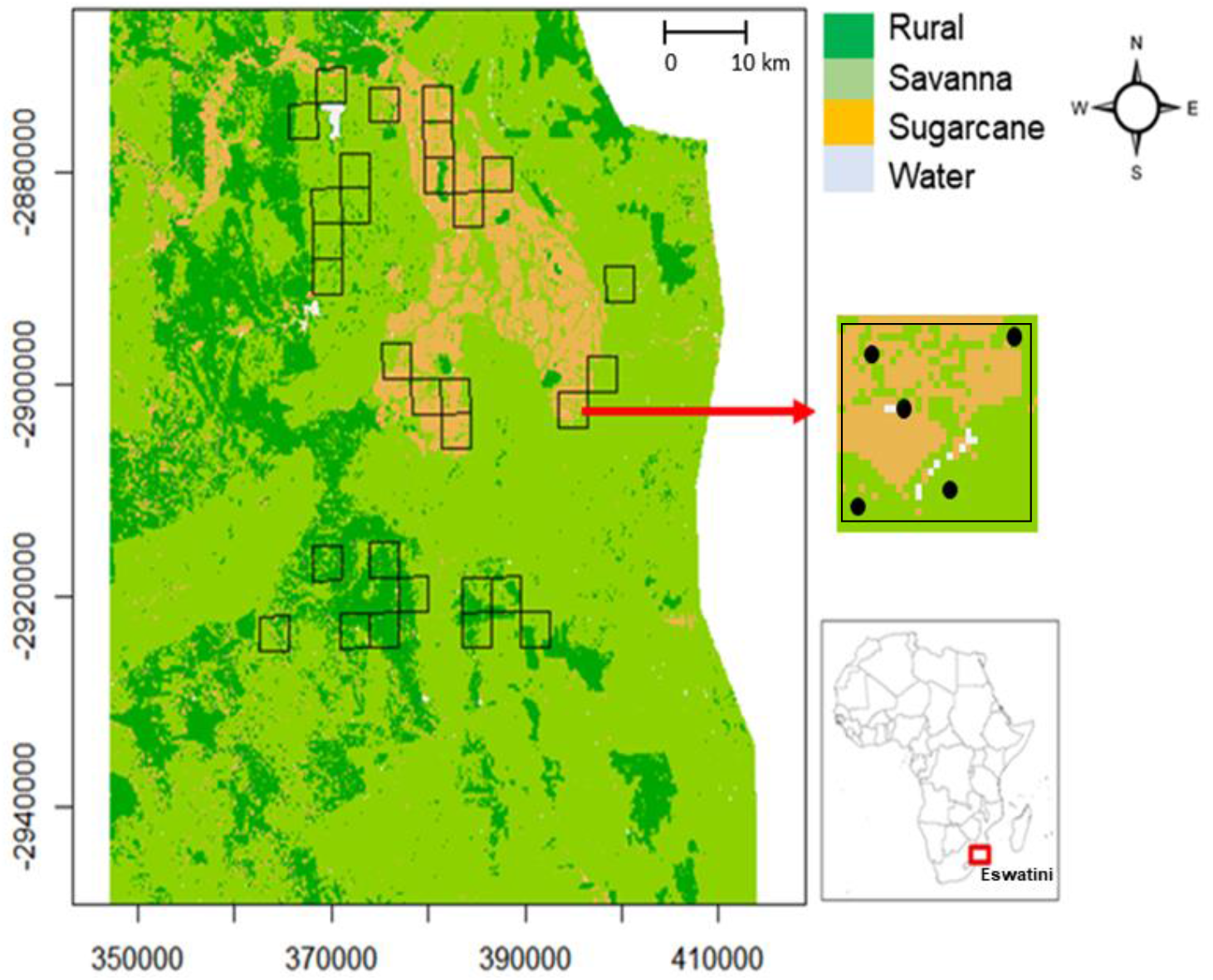
Map of the study region with sampling blocks outlined in black. The inset shows a close-up of one block, with Anabat points indicated by the black circles.

### 2.2 Land-Cover Classification

Land cover at our site is savanna vegetation (open savanna and woodland), commercial sugarcane plantations, and rural settlements, which included buildings, subsistence crops (primarily maize) and pasture for domestic livestock (Bailey et al., 2015; Monadjem and Reside, 2008). Several perennial rivers run through the study area and a number of dams occur here, mostly acting as reservoirs for the commercial plantations. Therefore, we classified land cover across the study region into four categories: rural settlements (hereafter “rural”), savannas, sugarcane plantations (hereafter “sugarcane”), and water. We used these four categories to create a classified raster of the region. First we carried out supervised classification in Google Earth Engine (www.earthengine.google.com) using a cloud-free Landsat 8 8-day raw composite image from March 21 – 29, 2016 at 30 m resolution. We then trained a voting support vector machine (voting SVM) classifier using 193 manually drawn polygons including each of the four land-cover categories. Resampling of the classified raster yielded an overall validity of 99.97%.

Because the rural land-cover class included crops and pasture that may have a similar spectral signature to savanna vegetation (Prestele et al., 2016), we incorporated population density to further distinguish rural areas from savanna. We used the population count raster for Eswatini from WorldPop projected for 2015 (WorldPop, 2013) to identify rural areas (Linard et al., 2012). We resampled this population count raster to the resolution of the classified raster using the nearest-neighbor algorithm. We overlaid the population raster on the classified raster and reclassified any cells with population count >1 as rural (Figure 1).

### 2.3 Acoustic Sampling

To capture variation in landscape cover across our study site we created a grid of 3 km^2^ (~1.73 km × ~1.73 km) blocks (hereafter “block”). We then overlaid this grid on the classified raster. We randomly selected 30 blocks (out of a possible 780) for acoustic surveys. These blocks were stratified between the three land-cover categories, with ten blocks for each type (10 rural, 10 savanna, 10 sugarcane). Within each block, we deployed five Anabat Express detectors (Titley, Inc., Ballina, Australia) at randomly placed points (hereafter “points”) from November 2015 – July 2016 (Figure 1). Each detector was attached to a tree trunk or electric pole at 1.5 m above the ground. Anabat detectors were set to record starting half an hour before sunset and continued recording for six hours. Each block was surveyed twice per season (wet: November – March; dry: May – July) for a total of four survey nights.

### 2.4 Classification of Bat Calls

We first trained a support vector machine (SVM) algorithm to classify bat calls based on calls from hand-released bats in the region (Monadjem et al., 2017). Five bat species (*Mops midas, Neoromicia nana, Scotophilus dinganii, Miniopterus natalensis*, and *Hipposideros caffer*) have calls that are distinctive and do not overlap in parameters with other species in the region. These species could be individually identified by the SVM algorithm. Several other species exhibit overlap in their call parameters (Monadjem et al., 2017) and were therefore grouped together into the following three “sono-species” during classification:

1. *Chaerephon pumilus – Mops condylurus – Taphozous mauritianus*
2. *Neoromicia zuluensis – Nycticeinops schlieffeni – Pipstrellus hesperidus – Scotophilus viridis*
3. *Rhinolophus blasii – R. darlingi – R. simulator*

In addition, we manually searched through bat files to identify calls from the two *Myotis* species from the region (*Myotis bocagii* and *M. tricolor*), which are visually distinctive from other bat species in the region, but have highly variable call parameters (Monadjem et al., 2017).

We examined the echolocation calls recorded at each point with the program ANALOOK (Chris Corben, version 4.8, http://www.hoarybat.com). Calls were first filtered to remove files with only noise and no bat calls. We then extracted the call parameters from those Anabat files that passed the noise filter. These parameters describe each bat pulse within a pass. The SVM algorithm classified bat calls at the level of the bat pulse within a pass. In order to be counted, four consecutive pulses had to be classified as the same sono-species. We validated the classifier by comparing a manual identification to the SVM classifier for 639 calls. SVM classification and manual identification were in agreement for 98.3% of the 639 validation calls.

We standardized the number of calls per sono-species by counting each species a maximum of once per minute (Miller, 2001). Finally, we grouped classified calls from each species or species group into three foraging guilds based on their wing morphology, echolocation, and foraging ecology: aerial foragers, edge foragers, and clutter foragers (Arita and Fenton, 1997; Meyer et al., 2004; Monadjem et al., 2010; Monadjem and Reside, 2008; Schnitzler and Kalko, 2001). Aerial foragers are adapted to fast, less maneuverable flight in open areas, while clutter foragers are adapted to slower, more maneuverable flight within dense vegetation; edge foragers are intermediate in terms of flight speed and maneuverability and often use vegetation at the edge of more open areas (Arita and Fenton, 1997; Meyer et al., 2004; Monadjem et al., 2010; Monadjem and Reside, 2008; Schnitzler and Kalko, 2001) (Table 1).

**Table 1.** Definition of foraging guilds and classification of bat species by foraging guild

| Foraging guild | Wing morphology | Echolocation | Foraging ecology | Species / Species Group |
| --- | --- | --- | --- | --- |
| Aerial | Long and narrow, high wing-loading | Low duty-cycle - Quasi-constant frequency | Open spaces, high altitudes | *Chaerephon pumilus* – *Mops condylurus* – *Taphozous mauritanus* group<br>*Mops midas* |
| Edge | Intermediate length, width, and wing loading | Low duty-cycle frequency-modulated or frequency-modulated-quasi-constant frequency | Edges of dense vegetation | *Neoromicia nana*<br>*Scotophilus dinganii*<br>*Neoromicia zuluensis* – *Nycticeinops schlieffeni* – *Pipstrellus hesperidus* – *Scotophilus viridis* group<br>*Myotis bocagii* – *Myotis tricolor* group<br>*Miniopterus natalensis* |
| Clutter | Short and broad, low wing-loading | Constant frequency | Dense, cluttered vegetation | *Rhinolophus blasii* – *R. darlingi* – *R. simulator* group |

### 2.5 Fine- and Landscape-Scale Metrics

We quantified the environment at two spatial scales: a fine scale around each sampling point and the landscape scale within each sampling block. At the fine scale, we measured vegetation cover and structure. In order to do so, we established a 30 m transect in each of the cardinal directions from the sampling point. We evaluated canopy and ground cover at the sampling point where the Anabat detector was placed and at points at 10 m intervals along each 30 m transect (total of thirteen measurements) while shrub cover was measured along the length of each 10 m interval within each transect (total of twelve measures). We measured the canopy cover using a spherical densiometer (Forestry Suppliers, Inc., Jackson MS) (Lemmon, 1956). We visually estimated ground cover in 1 × 1 m quadrats. We classified ground cover as: sugarcane, crop (all crops other than sugarcane), grass, bare ground, and water. We measured shrub cover, woody vegetation <2 m in height (Edwards, 1983), using the line intercept method (Canfield, 1941). For each sampling point, we took the mean canopy cover and ground cover from the thirteen points where we took these measures and the mean shrub cover from the twelve transects around the sampling point. We also measured the distance from each Anabat sampling point to the nearest water source because bats are known to use and forage around water bodies and riparian corridors (Monadjem and Reside, 2008; Pinto and Keitt, 2008; Sirami et al., 2013), using the function “gDistance” in the package rgeos (Bivand et al., 2017).

We calculated a variety of land-cover composition and configuration metrics within each sampling block (Gustafson, 1998). To account for land-cover composition, we measured the percent cover of savanna, rural, sugarcane, and water. For configuration metrics, we used savanna edge density because many bats use edges of natural vegetation (Chambers et al., 2016; Ethier and Fahrig, 2011; Mendes et al., 2017; Müller et al., 2012) and the savanna splitting index (hereafter “savanna splitting”), to account for the connectivity of savanna land cover, which may also be important for bats (Frey-Ehrenbold et al., 2013). We calculated all land-cover composition and configuration metrics using the “ClassStat” function in the SDMTools package (VanDerWal et al., 2014) in R version 3.3.3 (R Core Team, 2013).

We calculated pairwise correlations between all fine-scale metrics and all landscape-scale metrics using the function “rcorr” in the package Hmisc (Harrell, 2006). We found no correlations >0.7 among either the fine- or landscape-scale metrics that we used in our models.

### 2.6 Statistical Analysis

#### 2.6.1 Bat activity

We measured the response of aerial, edge, and clutter foragers’ activity at two scales: fine scale and landscape scale. At the fine scale, we summed the total number of calls at each Anabat point over all the sampling nights per season. For the landscape scale, we summed the number of bat calls per season from all Anabat detectors within the block. We measured bat response separately for each season (wet vs. dry) at both spatial scales because levels of bat activity are known to vary between seasons due to changes in temperature, precipitation, prey abundance and water availability (Cisneros et al., 2015; Ferreira et al., 2017; Klingbeil and Willig, 2010; Mendes et al., 2014).

We evaluated *a priori* suites of models to explain bat activity at both the fine and landscape scales. Each fine scale model included one of the fine-scale measures of vegetation structure: canopy cover, shrub cover, sugarcane cover, bare ground cover, water cover, and distance to water. We also included a null model (Table 2). To evaluate these models, we used generalized linear mixed models with the function “glmer” in the package lme4 (Bates et al., 2015), with a Poisson distribution to measure the response to fine-scale covariates. We used an offset term to account for the different number of sampling nights per point (Warton et al., 2015), due to occasional equipment failure. We used “block” as a random effect in order to account for spatial autocorrelation between points within the same block (Bailey et al., 2017).

**Table 2.**
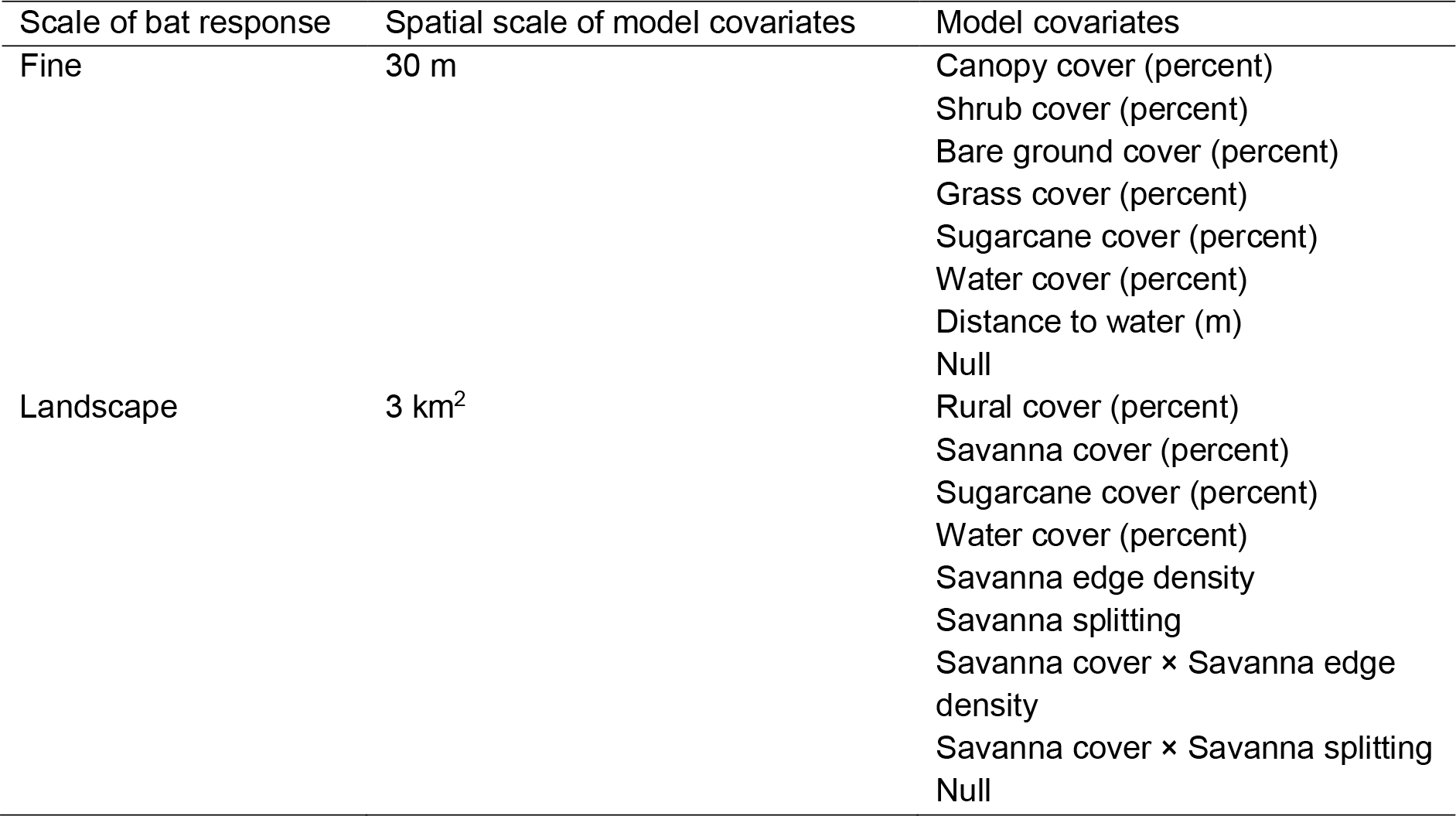
List of models used for each spatial scale. “×” indicates interactive term in models.

Landscape-scale models included one measure of landscape composition or configuration: rural, sugarcane, savanna, and water cover, edge density of savanna, or savanna splitting index. We also included two models with interactive effects between savanna composition and configuration: savanna cover × savanna edge density and savanna cover × savanna splitting (Table 2). We included interaction terms in order to determine whether savanna configuration may exacerbate or mitigate the effects of reduced savanna cover (composition). We used generalized linear models in base R v. 3.3.3 (www.r-project.org) with a Poisson distribution to measure the response to covariates at the landscape scale. Because the landscape response was aggregated at the block level, we did not include a random term to account for block. We used an offset term that was the sum of the number of sampling nights from all detectors within the block (Warton et al., 2015).

Within each scale and for each season, we compared models using Akaike Information Criterion corrected for small sample size (AICc) using the function “model.sel” in the package MuMIn (Barton, 2017). We considered models within 2 AICc units to be competing models. We then compared the point response models to each other and the best block response models to each other, using AICc. We evaluated the parameters of the top models by examining their 95% Confidence Intervals (CIs) and considered those that did not cross 0 to be relevant. We then graphed relevant parameters to understand how activity changes across variables of interest.

Finally, we compared the fit of the overall best fine-scale models to the overall best landscape-scale models using Pseudo *R^2^* (McFadden, 1974). Pseudo *R^2^* measures the deviance explained by a given model compared to the null model. We used Pseudo *R^2^* because the local and landscape models had different responses (e.g. activity at Anabat points vs. activity summed across all Anabat points within a block, respectively) and are therefore not directly comparable.

## 3. Results

We recorded acoustic data for a total of 3,408 hours during 120 sampling nights across the 30 sampling blocks. During this period, we identified a total of 69,897 bat calls. These calls were predominantly from aerial bats (n=48,466), followed by edge bats (n=21,361), and finally clutter bats (n=70). In general, we found that all three guilds responded more to the landscape scale than the fine scale and this response was stronger in the dry season than the wet season, but each guild responded differently to the landscape (Table 3).

**Table 3.** Top model and any competing models for each guild at each spatial scale in each season. See Supplemental Information for full model selection tables.

| Guild | Scale | Season | Top Model | Pseudo $R^2$ | Competing models |
| --- | --- | --- | --- | --- | --- |
| Aerial | Fine | Wet | Water cover | 0.04 | None |
|  |  | Dry | Water cover | 0.07 | None |
|  | Landscape | Wet | Savanna cover × Savanna splitting | 0.12 | None |
|  |  | Dry | Savanna cover × Savanna splitting | 0.28 | None |
| Edge | Fine | Wet | Shrub cover | 0.04 | None |
|  |  | Dry | Distance to water | 0.08 | None |
|  | Landscape | Wet | Savanna cover × Savanna splitting | 0.09 | None |
|  |  | Dry | Savanna cover × Savanna splitting | 0.18 | None |
| Clutter | Fine | Wet | Grass cover | 0.04 | Bare ground cover<br>Null model |
|  |  | Dry | Sugarcane cover | 0.04 | Water cover |
|  | Landscape | Wet | Rural cover | 0.21 | None |
|  |  | Dry | Water cover | 0.48 | None |

### 3.1 Aerial foraging guild

At the fine scale, the best model to explain activity of aerial foragers during both seasons was water cover. Activity increased with increasing water cover during both the wet season (β = 0.09, [95% confidence interval: 0.08, 0.10]) and dry season (β = 0.14 [0.13, 0.16]). There were no other competing models (Table 3, Table S1, Fig. 2). The Pseudo *R^2^* for top models in both seasons was relatively low, though higher in the dry (0.07 vs. 0.04) (Table 3).

**Figure 2.**
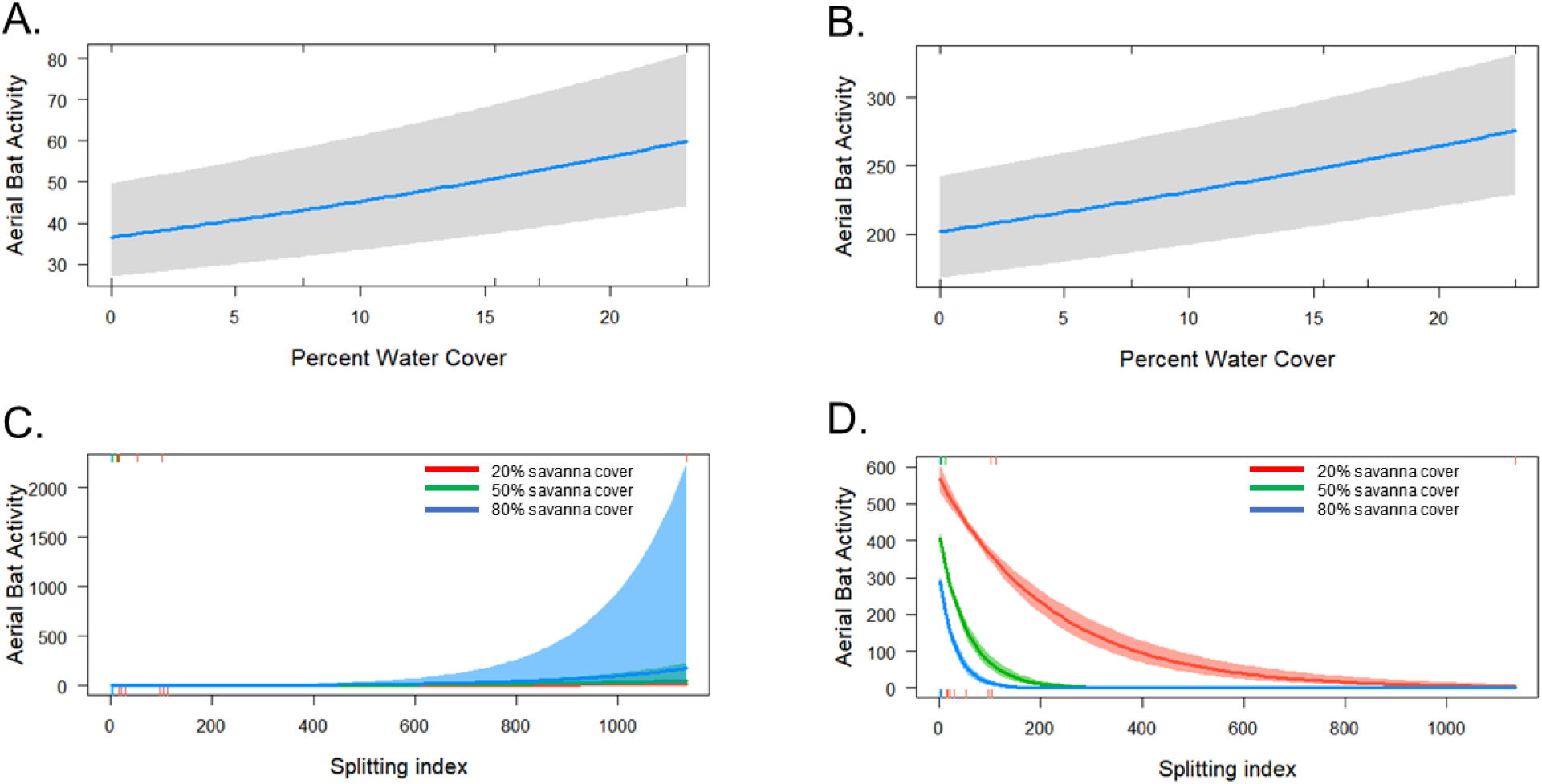
Response of aerial foraging guild bats at the fine scale in the A. wet season, B. dry season and at the landscape scale in the C. wet season and D. dry season.

At the landscape scale, the best model to explain activity in both seasons was a model with interactive effects of savanna cover and savanna splitting (Table 3). There was a positive relationship with activity in the wet season (savanna cover: β = 0.09 [0.03, 0.15]; savanna splitting: β = 0.66 [0.36,0.95]; interaction: β = 0.19 [0.02,0.36]) and negative relationship in the dry season ([−1.13, −0.87]; savanna splitting: β = −3.97 [−4.63, −3.31]; interaction: β = −2.47 [-2.85, −2.08]). During the wet season, activity increased more quickly with increasing savanna splitting where there was greater savanna cover. In the contrast, in the dry season, activity decreased with increasing savanna splitting, with a more rapid decline when savanna cover was higher (Fig. 2). There were no other competing models (Table 3, Table S1). Pseudo *R^2^* was over twice as high for dry season models as the wet (0.28 vs. 0.12) (Table 3).

### 3.2 Edge foraging guild

At the fine scale, the best model explaining activity of edge bats during the wet season was a model with percent shrub cover. Shrub cover was a relevant predictor of bat activity, which decreased with increasing cover (β = −0.23 [−0.25, −0.20]). The best model to explain bat activity in the dry season was a model with distance to water. Bat activity increased with decreasing distance from water (β = −0.77 [−0.88, −0.67]). There were no other competing models to explain edge bat activity during either season (Table 3, Table S1). Similar to aerial bats, Pseudo *R^2^* was twice as high for dry season models as the wet (0.08 vs. 0.04) (Table 3, Table S2, Figure 3).

**Figure 3.**
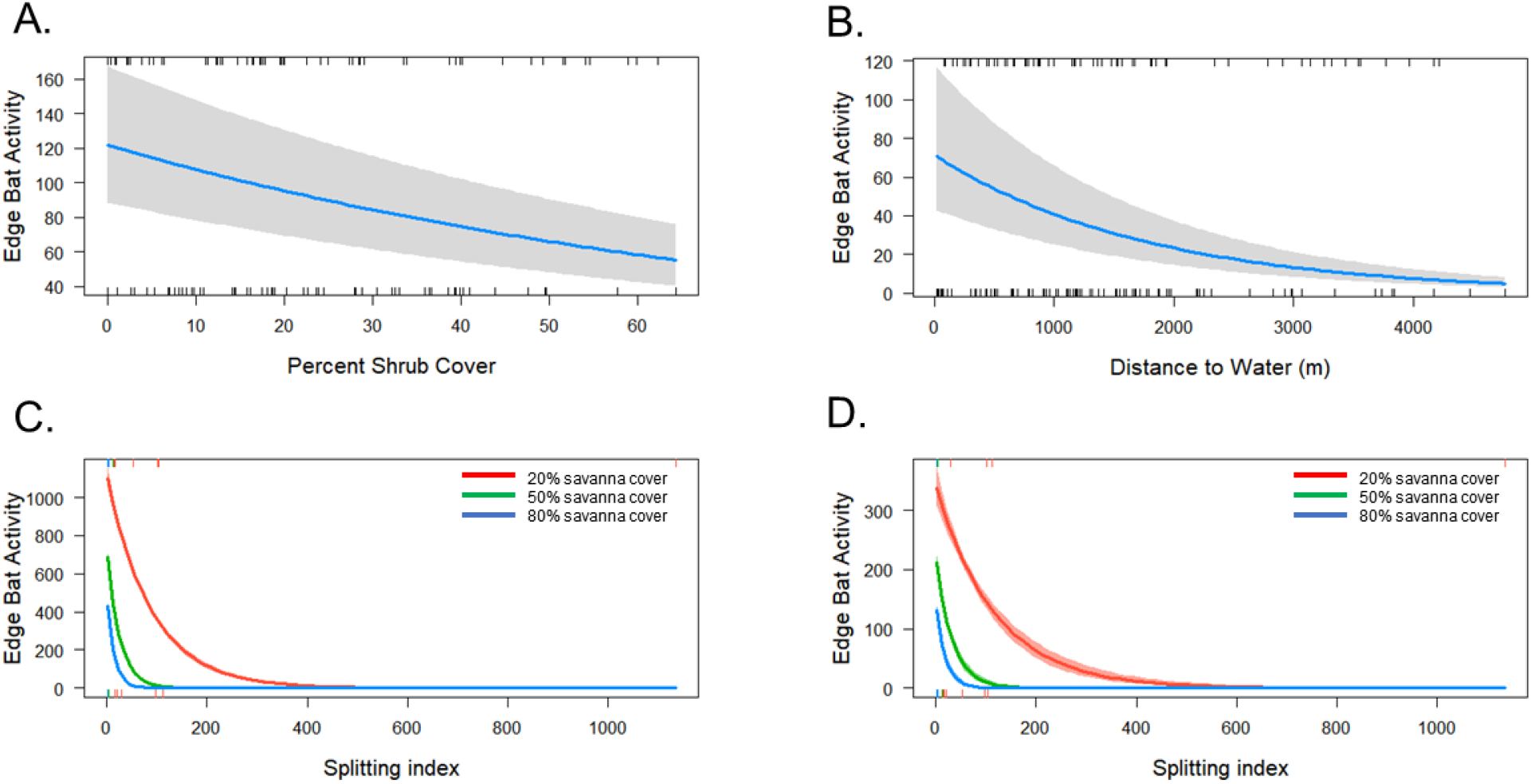
Response of edge foraging guild bats at the fine scale in the A. wet season, B. dry season and at the landscape scale in the C. wet season and D. dry season.

At the landscape scale, the best model to explain the activity of edge bats was a model with the interaction between savanna cover and splitting (Table 3, Table S1). The response was similar in both seasons, activity decreased with splitting and cover (wet: savanna cover: β = −1.87 [−1.98, − 1.76]; savanna splitting: β = −8.6 [−9.16, −8.05]; dry: savanna cover: β = −1.57 [−1.78, −1.36]; savanna splitting: β = −6.73 [−7.78, −5.70]). However, the decrease in bat activity with savanna splitting was reduced on blocks with less savanna (wet: interaction: β = −5.1 [−5.42, −4.78]; dry; interaction: β = −4.05 [−4.66, −3.45]) (Fig. 3). There were no competing models (Table 3, Table S1). The dry season model (Pseudo *R*^2^=0.18) fit the data better than the wet season model (Pseudo *R*^2^=0.09). (Table 3).

### 3.3 Clutter foraging guild

The best model of activity of clutter bats at the fine scale in the wet season was a model with the variable grass cover, but with a 95% CI that included 0 and hence it was not a relevant predictor (β = 0.50 [−0.04, 1.07]). The null model and a model with bare ground cover were also competing models but bare ground was also not a relevant predictor (β = −0.50 [−1.31, 0.17]) In the dry season the best model included the variable sugarcane cover, which was a relevant predictor (β = 0.36 [0.09, 0.62]); bat activity increased with increasing sugarcane cover. A model including the variable water cover was also a competing model, but it was not a relevant predictor (β = −0.53 [−0.71, 0.55]) (Table 3, Table S3, Figure 4). The model fit for the top models in both seasons was comparable (Pseudo *R*^2^=0.04).

**Figure 4.**
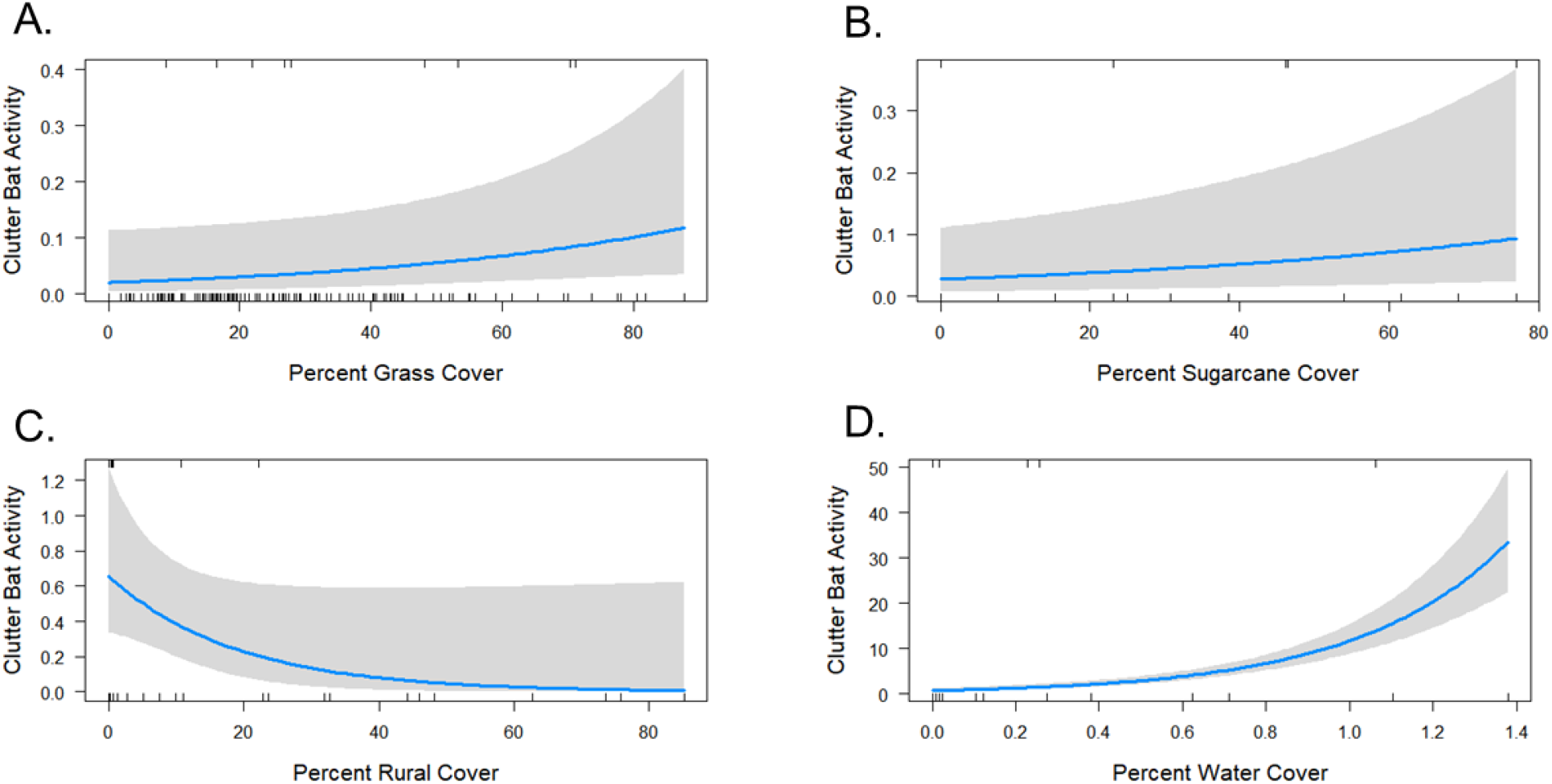
Response of clutter foraging guild bats at A. wet season, B. dry season and at the landscape scale in the C. wet season and D. dry season.

At the landscape scale, the best model to explain the activity of clutter bats during the wet season was the amount of rural land cover. Bat activity decreased as the amount of rural land in a block increased (β = −1.36 [−3.37, −0.32]). During the dry season the best model explaining bat activity was water cover, with activity increasing with increasing water (β = 1.03 [0.85, 1.22]). There were no competing models at the landscape scale in either season. The top dry season model for clutter foragers (Pseudo *R*^2^=0.48) fit data better than our top wet season model (Pseudo *R*^2^=0.21) (Table 3, Table S3, Figure 4).

## 4. Discussion

This study demonstrates the role of both fine-scale vegetation structure and landscape-scale composition and configuration in shaping bat activity within a savanna undergoing rapid land-use and land-cover change (Bailey et al., 2015). Across all three bat foraging guilds, we found that activity was best explained by landscape-scale characteristics rather than fine-scale vegetation parameters. Previous studies have reported that bats with larger home ranges respond more strongly to broad-scale features of the landscape, while bats with smaller home ranges respond more to fine-scale vegetation structure (Ferreira et al., 2017; Fuentes-Montemayor et al., 2013; Klingbeil and Willig, 2010; Pinto and Keitt, 2008). Although clutter bats have much smaller home ranges than edge or aerial bats, they may still fly up to 2 km per night, which may explain the relevance of broader scale landscape features as reported here and elsewhere (Fenton, 1990; Fenton and Rautenbach, 1986; Monadjem et al., 2009). Our results suggest any conservation planning or assessment of bat biodiversity in tropical African savannas should consider land cover at broad scale (>3 km^2^). The use of inappropriate spatial scales may limit the effectiveness of conservation actions or mitigation measures. Indeed, there is evidence that mitigations (such as agro-environmental measures) that are implemented only at fine scales, such as leaving hedgerows or small patches of natural vegetation, may be ineffective in promoting or maintaining bat activity (Fuentes-Montemayor et al., 2011).

We found that these landscape characteristics explained more of the bat activity response in the dry season than the wet season for all three foraging guilds. Seasonal responses in bat activity are common and have been found in tropical savannas of this region (Mtsetfwa et al., 2018; Taylor et al., 2013) as well as other parts of the world (Cisneros et al., 2015; Ferreira et al., 2017; Klingbeil and Willig, 2010; Mendes et al., 2014). During the wet season, essential resources, such as insect prey and water, are more abundant (Fukui et al., 2006; Hagen and Sabo, 2012; Salsamendi et al., 2012) and therefore bats might be less constrained or affected by landscape composition and configuration. The effect of landscape may be more pronounced in the dry season because resources, particularly water, become scarce (Korine et al., 2016).

While we predicted that bats would respond more strongly to landscape composition than configuration, we found that both composition and configuration, particularly fragmentation of savanna land cover, were important for aerial and edge foraging bats. Bats have been shown to exhibit both negative and positive responses to fragmentation; these responses are often species- or guild-specific (Cosson et al., 1999; Estrada-Villegas et al., 2010; Ethier and Fahrig, 2011; Meyer et al., 2016). While some studies have found that the amount of natural cover is more important than fragmentation for bats (Meyer and Kalko, 2008), here we find an interactive effect between savanna cover and fragmentation. This interaction suggests that the effect fragmentation has on aerial and edge foraging bats depends on the amount of savanna cover. When savanna cover is high (>50%), fragmentation results in a steep decline in bat activity. At lower savanna cover (20%), fragmentation still has a negative effect, but the reduction in bat activity is less pronounced, perhaps because at this level the remaining savanna essentially exists in small fragments only. Alternatively, the decline in bat activity may be less pronounced at lower savanna cover because the other land-cover types (e.g. sugarcane and rural) in the landscape provide adequate resources, such as food, water, or roost sites.

Bat activity tends to increase in lower intensity agricultural systems, such as agroforestry and organic farms, at least in the few studies that have investigated this relationship (Cleary et al., 2016; Park, 2015; Wickramasinghe et al., 2003). However, we found that clutter bats responded negatively to rural cover, which is comprised of low-intensity small-holder crops, homes, pasture and dirt roads. These areas are typically very open, with large areas of bare ground and few trees or shrubs. The lack of dense vegetation likely limits the ability of clutter forager bats to use rural areas (Monadjem and Reside, 2008; Schnitzler and Kalko, 2001). On the other hand, we found that sugarcane had a significant, positive effect on clutter bats at the fine scale in the dry season. During this season, sugarcane plantations may offer resources, such as water from dams or irrigation canals and insects that are scarce in savannas or rural areas. In addition, sugarcane is densely planted and may reach two meters in height and therefore may provide suitable habitat for clutter foragers. The resemblance of vegetation structure to native vegetation in areas of agricultural land use may be more important for bats than the production intensity.

We found that water was important for all three foraging guilds in the dry season, although there was variation in the spatial scale at which water drove activity for each guild. Water availability is important for bats in general, providing both water for drinking and insect foraging (Adams, 2010; Adams and Hayes, 2008; Monadjem and Reside, 2008; Sherwin et al., 2013; Sirami et al., 2013). Water may play an even more important role in savannas, where availability might be lower than other tropical biomes, especially during dry seasons (Korine et al., 2016), and may drive bat movement and activity across the landscape (Geluso and Geluso, 2012; Rainho and Palmeirim, 2011). Because savannas, especially in arid and semi-arid areas, are at risk of future droughts and desertification (Engelbrecht et al., 2015; Stringer et al., 2009), water will likely become increasingly scarce for bats. Artificial water sources which are available year-round, such as the dams and canals within commercial agriculture areas and some villages, may provide an especially important resource for bats in this human-altered landscape (Sirami et al., 2013).

There are some limitations to the use of acoustic monitoring in this study. A number of echolocating species found in the region, such as *Nycteris thebaica* and *Kerivoula lanosa* cannot be detected by our acoustic detectors (Monadjem et al., 2017). Similarly, non-echolocating species such as the fruit bat *Epomophorus wahlbergi* (Shapiro and Monadjem, 2016) could also not be included. In addition, many species in the region cannot be distinguished from acoustic calls alone due to similarity in call parameters (Monadjem et al., 2017). While we see clear patterns by foraging guild, there could also be species-specific responses within guilds (Ethier and Fahrig, 2011; Fuentes-Montemayor et al., 2011; Gorresen et al., 2005; Gorresen and Willig, 2004; Pinto and Keitt, 2008), which we were unable to take into account.

Increasing levels of anthropogenic land-cover change around the world are cause for concern for many wildlife species and biodiversity as a whole (Foley et al., 2005; Jetz et al., 2007; Venter et al., 2016), including those in savannas (Laurance et al., 2014; Parr et al., 2014). However, despite the pressures of land-cover and land-use change, it is possible to conserve bats, and the ecosystem services they provide (Kunz et al., 2011; Taylor et al., 2018), in these changing savanna landscapes. Bats in savannas have a complex relationship with the landscape that varies by guild, season, and spatial scale. Therefore, any conservation or management strategies for bats in tropical savannas should consider the landscape at large scales (≥ 3 km), minimize fragmentation of existing savanna, especially in areas of high remaining coverage (>50%), and maintain water sources, both natural and artificial. Doing so can promote activity of aerial, edge, and clutter foragers across spatial and temporal scales.

## Acknowledgements

We thank Hervé Echecolonea for assistance in the field and Mandla Motsa, Smart Shabangu, Tal Fineberg, Isaac Magagula, Thea Litschka-Koen, Clifton Koen, Nick Jackson, Stephen Potts, Alan Howland and Kim Roques and All Out Africa, for help with logistics.

