## Supplemental Tables for "Response of Bat Activity to Land-Cover and Land-Use Change in Savannas is Scale-, Season-, and Guild-Specific"

**Supporting Information**

Table S1. Model selection table for aerial foraging guild bats indicating spatial scale, season, model names, degrees of freedom (df), loglikelihood, Akaike Information Criterion adjusted for small sample size (AICc), delta AICc (ΔAICc), and model weight.

| **Scale** | **Season** | **Model** | **df** | **Log- Likelihood** | **AICc** | **ΔAICc** | **Model weight** |
| --- | --- | --- | --- | --- | --- | --- | --- |
| Fine | Wet | Water cover | 3 | -5114.0 | 10234.1 | 0.0 | 1.0 |
|  |  | Shrub cover | 3 | -5185.9 | 10377.9 | 143.7 | 0.0 |
|  |  | Sugarcane cover | 3 | -5191.6 | 10389.4 | 155.3 | 0.0 |
|  |  | Grass cover | 3 | -5211.4 | 10428.9 | 194.8 | 0.0 |
|  |  | Bare ground cover | 3 | -5233.0 | 10472.1 | 238.0 | 0.0 |
|  |  | Distance to water | 3 | -5236.1 | 10478.3 | 244.2 | 0.0 |
|  |  | Canopy cover | 3 | -5265.2 | 10536.6 | 302.5 | 0.0 |
|  |  | Null | 2 | -5279.2 | 10562.4 | 328.3 | 0.0 |
|  | Dry | Water cover | 3 | -1873.6 | 3753.5 | 0.0 | 1.0 |
|  |  | Sugarcane cover | 3 | -1918.9 | 3844.1 | 90.6 | 0.0 |
|  |  | Grass cover | 3 | -1963.8 | 3933.8 | 180.3 | 0.0 |
|  |  | Bare ground cover | 3 | -1968.5 | 3943.2 | 189.8 | 0.0 |
|  |  | Null | 2 | -1976.4 | 3956.8 | 203.4 | 0.0 |
|  |  | Canopy cover | 3 | -1976.2 | 3958.5 | 205.1 | 0.0 |
|  |  | Distance to water | 3 | -1976.2 | 3958.6 | 205.2 | 0.0 |
|  |  | Shrub cover | 3 | -1976.3 | 3958.7 | 205.3 | 0.0 |
| Landscape | Wet | Savana cover x Savanna splitting | 4 | -9785.2 | 19580.0 | 0.0 | 1.0 |
|  |  | Savanna splitting | 2 | -9796.1 | 19596.6 | 16.6 | 0.0 |
|  |  | Water cover | 2 | -10889.2 | 21782.9 | 2203.0 | 0.0 |
|  |  | Savanna cover x Edge density | 4 | -10915.1 | 21839.8 | 2259.9 | 0.0 |
|  |  | Savanna cover | 2 | -11027.2 | 22058.9 | 2478.9 | 0.0 |
|  |  | Sugarcane cover | 2 | -11027.2 | 22058.9 | 2478.9 | 0.0 |
|  |  | Rural cover | 2 | -11031.9 | 22068.3 | 2488.3 | 0.0 |
|  |  | Edge density | 2 | -11135.5 | 22275.4 | 2695.4 | 0.0 |
|  |  | Null | 1 | -11141.4 | 22284.9 | 2704.9 | 0.0 |
|  | Dry | Savana cover x Savanna splitting | 4 | -2338.8 | 4687.3 | 0.0 | 1.0 |
|  |  | Savanna splitting | 2 | -2501.4 | 5007.2 | 320.0 | 0.0 |
|  |  | Savanna cover x Edge density | 4 | -2556.0 | 5121.5 | 434.3 | 0.0 |
|  |  | Water cover | 2 | -2737.9 | 5480.2 | 792.9 | 0.0 |
|  |  | Savanna cover | 2 | -2870.3 | 5745.1 | 1057.9 | 0.0 |
|  |  | Sugarcane cover | 2 | -2870.3 | 5745.1 | 1057.9 | 0.0 |
|  |  | Edge density | 2 | -3046.0 | 6096.4 | 1409.1 | 0.0 |
|  |  | Rural cover | 2 | -3217.4 | 6439.2 | 1751.9 | 0.0 |
|  |  | Null | 1 | -3224.4 | 6451.0 | 1763.7 | 0.0 |

Table S2. Model selection for edge foraging guild bats indicating spatial scale, season, model names, degrees of freedom (df), loglikelihood, Akaike Information Criterion adjusted for small sample size (AICc), delta AICc (ΔAICc), and model weight.

| **Scale** | **Season** | **Model** | **df** | **Log-likelihood** | **AICc** | **ΔAIC** | **Model weight** |
| --- | --- | --- | --- | --- | --- | --- | --- |
| Fine | Wet | Shrub cover | 3 | -3911.5 | 7829.2 | 0.0 | 1.0 |
|  |  | Bare ground cover | 3 | -3988.3 | 7982.7 | 153.5 | 0.0 |
|  |  | Grass cover | 3 | -3999.3 | 8004.8 | 175.5 | 0.0 |
|  |  | Water cover | 3 | -4014.2 | 8034.5 | 205.3 | 0.0 |
|  |  | Sugarcane cover | 3 | -4028.3 | 8062.8 | 233.6 | 0.0 |
|  |  | Canopy cover | 3 | -4035.9 | 8077.9 | 248.7 | 0.0 |
|  |  | Null | 2 | -4037.6 | 8079.2 | 250.0 | 0.0 |
|  |  | Distance to water | 3 | -4037.2 | 8080.5 | 251.3 | 0.0 |
|  | Dry | Distance to water | 3 | -1804.0 | 3614.3 | 0.0 | 1.0 |
|  |  | Water cover | 3 | -1840.9 | 3687.9 | 73.6 | 0.0 |
|  |  | Canopy cover | 3 | -1880.9 | 3768.0 | 153.8 | 0.0 |
|  |  | Sugarcane cover | 3 | -1894.8 | 3795.7 | 181.5 | 0.0 |
|  |  | Bare ground cover | 3 | -1899.3 | 3804.9 | 190.6 | 0.0 |
|  |  | Grass cover | 3 | -1902.5 | 3811.3 | 197.0 | 0.0 |
|  |  | Null | 2 | -1918.4 | 3841.0 | 226.7 | 0.0 |
|  |  | Shrub cover | 3 | -1917.6 | 3841.3 | 227.0 | 0.0 |
| Landscape | Wet | Savanna cover x Savanna splitting | 4 | -8364.4 | 16738.3 | 0.0 | 1.0 |
|  |  | Savanna cover x Edge density | 4 | -8775.5 | 17560.5 | 822.2 | 0.0 |
|  |  | Rural cover | 2 | -8959.7 | 17923.8 | 1185.5 | 0.0 |
|  |  | Water cover | 2 | -8985.4 | 17975.2 | 1236.9 | 0.0 |
|  |  | Savanna cover | 2 | -8992.1 | 17988.6 | 1250.3 | 0.0 |
|  |  | Sugarcane cover | 2 | -8992.1 | 17988.6 | 1250.3 | 0.0 |
|  |  | Savanna splitting | 2 | -9025.0 | 18054.4 | 1316.1 | 0.0 |
|  |  | Edge density | 2 | -9096.4 | 18197.3 | 1459.0 | 0.0 |
|  |  | Null | 1 | -9137.2 | 18276.6 | 1538.3 | 0.0 |
|  | Dry | Savanna cover x Savanna splitting | 4 | -1775.0 | 3559.6 | 0.0 | 1.0 |
|  |  | Water cover | 2 | -1799.6 | 3603.7 | 44.2 | 0.0 |
|  |  | Savanna cover x Edge density | 4 | -1896.6 | 3802.8 | 243.2 | 0.0 |
|  |  | Savanna splitting | 2 | -1925.8 | 3856.0 | 296.5 | 0.0 |
|  |  | Savanna cover | 2 | -1995.3 | 3995.1 | 435.6 | 0.0 |
|  |  | Sugarcane cover | 2 | -1995.3 | 3995.1 | 435.6 | 0.0 |
|  |  | Edge density | 2 | -2138.7 | 4281.9 | 722.3 | 0.0 |
|  |  | Rural cover | 2 | -2152.5 | 4309.5 | 749.9 | 0.0 |
|  |  | Null | 1 | -2153.8 | 4309.8 | 750.3 | 0.0 |

Table S3. Model selection for clutter foraging guild bats indicating spatial scale, season, model names, degrees of freedom (df), loglikelihood, Akaike Information Criterion adjusted for small sample size (AICc), delta AICc (ΔAICc), and model weight.

| **Scale** | **Season** | **Model** | **df** | **Log-likelihood** | **AICc** | **ΔAIC** | **Model weight** |
| --- | --- | --- | --- | --- | --- | --- | --- |
| Fine | Wet | Grass cover | 3 | -38.0 | 82.2 | 0.0 | 0.32 |
|  |  | Bare ground cover | 3 | -38.6 | 83.4 | 1.2 | 0.17 |
|  |  | Null | 2 | -39.7 | 83.4 | 1.3 | 0.17 |
|  |  | Shrub cover | 3 | -39.3 | 84.8 | 2.6 | 0.09 |
|  |  | Distance to water | 3 | -39.5 | 85.1 | 3.0 | 0.07 |
|  |  | Canopy cover | 3 | -39.6 | 85.3 | 3.1 | 0.07 |
|  |  | Sugarcane cover | 3 | -39.6 | 85.4 | 3.3 | 0.06 |
|  |  | Water cover | 3 | -39.6 | 85.5 | 3.3 | 0.06 |
|  | Dry | Sugarcane cover | 3 | -105.2 | 216.6 | 0.0 | 0.42 |
|  |  | Water cover | 3 | -105.6 | 217.4 | 0.8 | 0.28 |
|  |  | Distance to water | 3 | -106.4 | 219.1 | 2.5 | 0.12 |
|  |  | Grass cover | 3 | -107.2 | 220.5 | 3.9 | 0.06 |
|  |  | Canopy cover | 3 | -107.3 | 220.8 | 4.2 | 0.05 |
|  |  | Null | 2 | -108.6 | 221.3 | 4.8 | 0.04 |
|  |  | Bare cover | 3 | -108.6 | 223.3 | 6.8 | 0.01 |
|  |  | Shrub cover | 3 | -108.6 | 223.4 | 6.8 | 0.01 |
| Landscape | Wet | Rural cover | 2 | -24.3 | 53.0 | 0.0 | 0.74 |
|  |  | Savanna splitting | 2 | -26.6 | 57.7 | 4.7 | 0.07 |
|  |  | Water cover | 2 | -26.7 | 57.9 | 5.0 | 0.06 |
|  |  | Null | 1 | -28.3 | 58.8 | 5.8 | 0.04 |
|  |  | Savanna cover | 2 | -27.6 | 59.6 | 6.7 | 0.03 |
|  |  | Sugarcane cover | 2 | -27.6 | 59.6 | 6.7 | 0.03 |
|  |  | Edge density | 2 | -27.8 | 60.1 | 7.1 | 0.02 |
|  |  | Savanna cover x Savanna splitting | 4 | -26.6 | 62.8 | 9.8 | 0.01 |
|  |  | Savanna cover x Edge density | 4 | -27.1 | 63.8 | 10.9 | 0.00 |
|  | Dry | Water cover | 2 | -75.5 | 155.4 | 0.0 | 1.0 |
|  |  | Edge density | 2 | -108.5 | 221.5 | 66.1 | 0.0 |
|  |  | Savanna cover x Edge density | 4 | -107.6 | 224.8 | 69.4 | 0.0 |
|  |  | Savanna density x Savanna splitting | 4 | -113.5 | 236.7 | 81.3 | 0.0 |
|  |  | Rural cover | 2 | -119.5 | 243.4 | 88.0 | 0.0 |
|  |  | Savanna splitting | 2 | -126.6 | 257.6 | 102.2 | 0.0 |
|  |  | Null | 1 | -130.7 | 263.6 | 108.2 | 0.0 |
|  |  | Savanna cover | 2 | -130.3 | 265.1 | 109.7 | 0.0 |
|  |  | Sugarcane cover | 2 | -130.3 | 265.1 | 109.7 | 0.0 |
